# Regional Differences Following Partial Salivary Gland Resection

**DOI:** 10.1101/713230

**Authors:** Kevin J. O’Keefe, Kara A. DeSantis, Amber L. Altrieth, Deirdre A. Nelson, Ed Zandro M. Taroc, Adam R. Stabell, Mary T. Pham, Melinda Larsen

## Abstract

Regenerative medicine aims to repair, replace, or restore function to tissues damaged by aging, disease, or injury. Partial organ resection is not only a common clinical approach in cancer therapy, it is also an experimental injury model used to examine mechanisms of regeneration and repair in organs. We performed a partial resection, or partial sialodenectomy, in the murine submandibular salivary gland (SMG) to establish a model for investigation of repair mechanisms in salivary glands (SGs). After partial sialoadenectomy we performed whole gland measurements over a period of 56 days and found that the gland reached its maximum size 14 days after injury. We used microarray analysis and immunohistochemistry to examine mRNA and protein changes in glands over time. Microarray analysis identified dynamic changes in the transcriptome three days after injury that were largely resolved by day 14. At the 3 day time point, we detected gene signatures for cell cycle regulation, inflammatory/repair response, and extracellular matrix remodeling in the partially resected glands. Using quantitative immunohistochemistry, we identified a transient proliferative response throughout the gland, in which both secretory epithelial and stromal cells expressed Ki67 that was detectable at day 3 and largely resolved by day 14. IHC also revealed that while most of the gland underwent a wound healing response that resolved by day 14, a small region of the gland showed an aberrant sustained fibrotic response characterized by increased levels of ECM deposition and sustained Ki67 levels in stromal cells. The partial submandibular salivary gland resection model provides an opportunity to examine a normal healing response and an aberrant fibrotic response within the same gland to uncover mechanisms that prevent wound healing and regeneration in mammals. Understanding regional differences in the wound healing responses may ultimately impact regenerative therapies for patients.

## Introduction

Salivary gland (SG) hypofunction can occur as a side effect of radiation treatment in head and neck cancer patients as well as in autoimmune diseases including Sjögren’s Syndrome (SjS). Reduced salivary function negatively impacts patient quality of life, resulting in the feeling of dry mouth (xerostomia), increased incidence of dental caries, oral fungal infections, and difficulty in eating, speaking, swallowing, and tasting (Lombaert et al. 2017). For salivary gland cancer patients, partial SG resection is a common clinical practice for removal of cancerous regions, leading to a decrease in saliva flow one year after partial resection (Ge et al. 2016).

Resection-based preclinical models exist for studying the regenerative capacity of many organs including liver (Nevzorova et al. 2015), lung (Cowan and Crystal 1975), and heart (Xiong and Hou 2016). Here we investigated the response of the murine SMG to partial resection after removing 40% of the distal tip of the gland. Using microarray analysis to profile the transcriptome and immunohistochemistry (IHC) to examine temporal-spatial protein expression patterns in response to resection. We found that the resected SMG exhibited a global wound healing response, with an insufficient epithelial proliferation response to regenerate the gland. Interestingly, we also identified a small interior region of gland that responded to injury with a sustained fibrotic response.

## Materials and Methods

This is a preclinical animal study that conforms to Animal Research: Reporting of In Vivo Experiments (ARRIVE) guidelines. For details on materials and methods, see the Appendix.

## Results

### Submandibular Salivary Glands Respond to Partial Gland Resection

We performed a partial sialoadenectomy on 12 week old C57Bl/6J mice. We surgically removed approximately 40% of the distal tip of the left SMG at a natural tissue margin (Fig. 1A), noting low variance in the size and weight of the tissue pieces removed (Fig. 1B, C). After partially resecting the glands we allowed the gland to recover for 3, 14, or 56 days. We measured the length (Fig. 1D) and width (Fig. 1E) of the resected and control glands from non-operated animals *in situ*, and removed glands to weigh them (Fig. 1F). Contralateral right glands in operated animals were also measured (Appendix Fig. 1). The resected glands appeared to respond to the injury showing measureable changes in gland length and width, with an increasing trend in gland weight.

**Figure 1.**
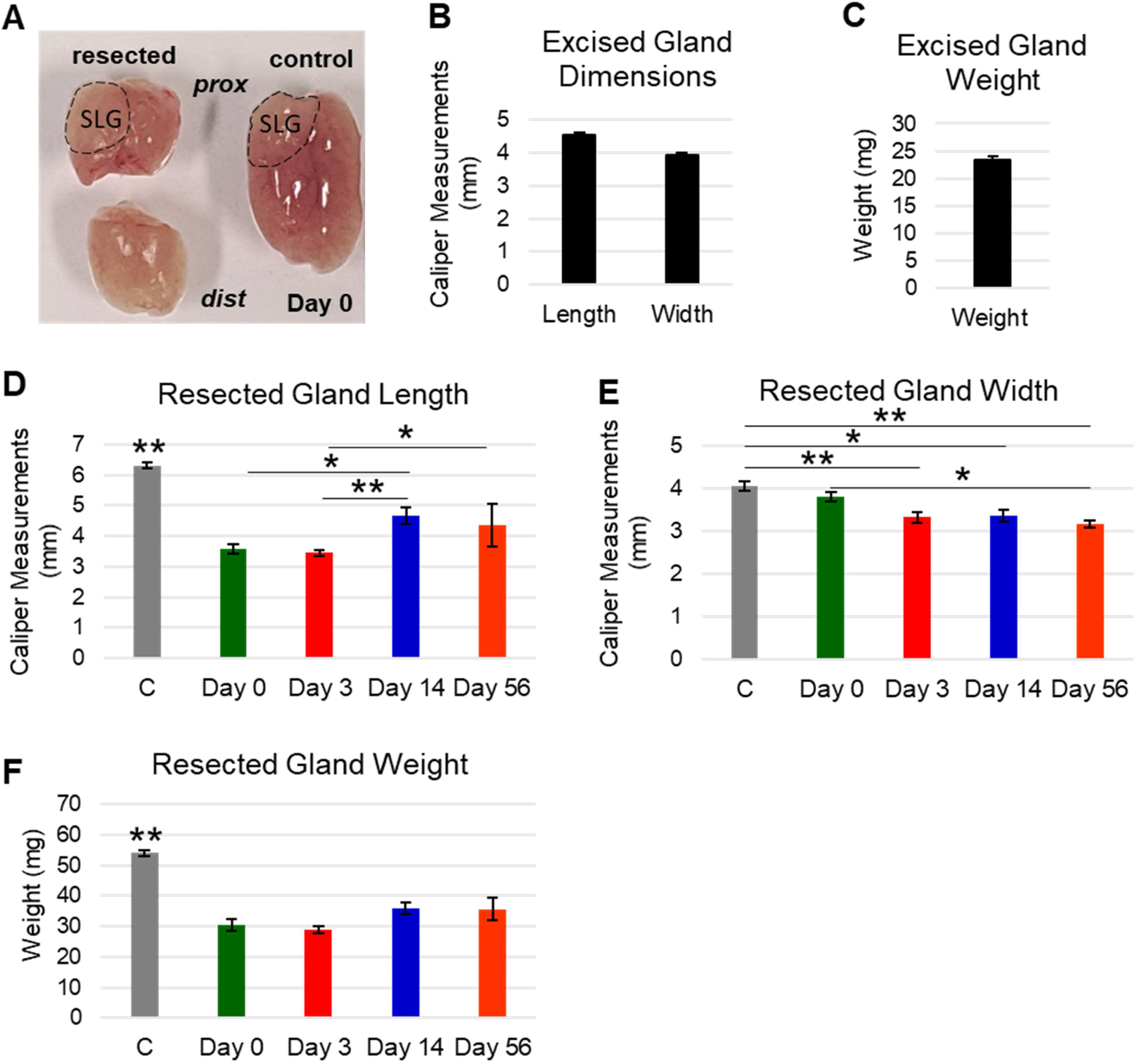
Submandibular Salivary Glands Respond After Partial Gland Resection. **A)** Image showing partially resected submandibular gland (top left) and excised gland piece (bottom left), compared to unmanipulated control gland (right) at day 0. Image shows proximal to distal gland orientation with the sublingual gland (SLG) indicated with black dotted line. **B)** Excised gland dimensions were measured with calipers in millimeters (mm). Length (n=32) and width (n=32) measurements show consistency in surgical technique. **C)** Excised gland weight milligrams (mg) (n=32). **D)** Gland length measured *in situ* in control (C) (n=5), resection with immediate tissue harvest (Day 0) (n=9), 3 days post resection (Day 3) (n=11), 14 days post resection (Day 14) (n=8) and 56 days post resection (Day 56) (n=4). Glands respond to surgery with an increase in length that is maximal at Day 14. **E)** Resected gland weight (mg) measured at all time points shows that glands do not accumulate increased mass after resection and do not reach the size of C glands. **F)** Resected gland widths (mm) show that glands get progressively narrow following partial gland resection. **G)** Weights of contralateral glands (mg) demonstrate that contralateral glands gain weight after resection and reach maximum at Day 14. Error bars are S.E.M. Statistical Tests: One-Way ANOVA Tukey’s HSD, * p≤0.05, ** p≤0.01.

### Molecular Profiling Reveals an Intact Wound Healing Response and Cell Cycle Entry

To gain insight into the nature of the gland response to resection at the level of RNA, we compared the transcriptome profile of the partially resected glands at day 3 and 14, in comparison to day 3 mock surgery controls. We examined the transcriptomes of each treatment using Clariom S microarrays, which include 20,000 well-annotated genes and assess only exons known to be expressed. Principle component analysis (PCA) revealed that the transcriptomes of the glands at day 3 were distinct from the other samples, whereas the transcriptomes of the day 14 glands more closely resembled that of the control glands (Fig. 2A). Filtering for genes changed more than 2-fold relative to control confirmed that there were many more genes that were differentially regulated at day 3 than at day 14 (Fig. 2B). Volcano plots revealed at 3 days there were more genes increased than were decreased (Fig. 2C), and these changes in gene expression were largely diminished after 14 days (Fig. 2D). We used Metascape (Zhou et al. 2019) to identify categories of genes that changed most significantly at days 3 after partial gland resection. Metascape clustering analysis revealed changes in genes involved in cell cycle and repair processes including, hemostasis, inflammation, cytokine production, and adaptive immunity (Fig. 2E and Appendix Fig. 3). These data indicate that the partially resected glands undergo changes in gene expression immediately after resection consistent with a wound healing response that is largely resolved at day 14.

**Figure 2.**
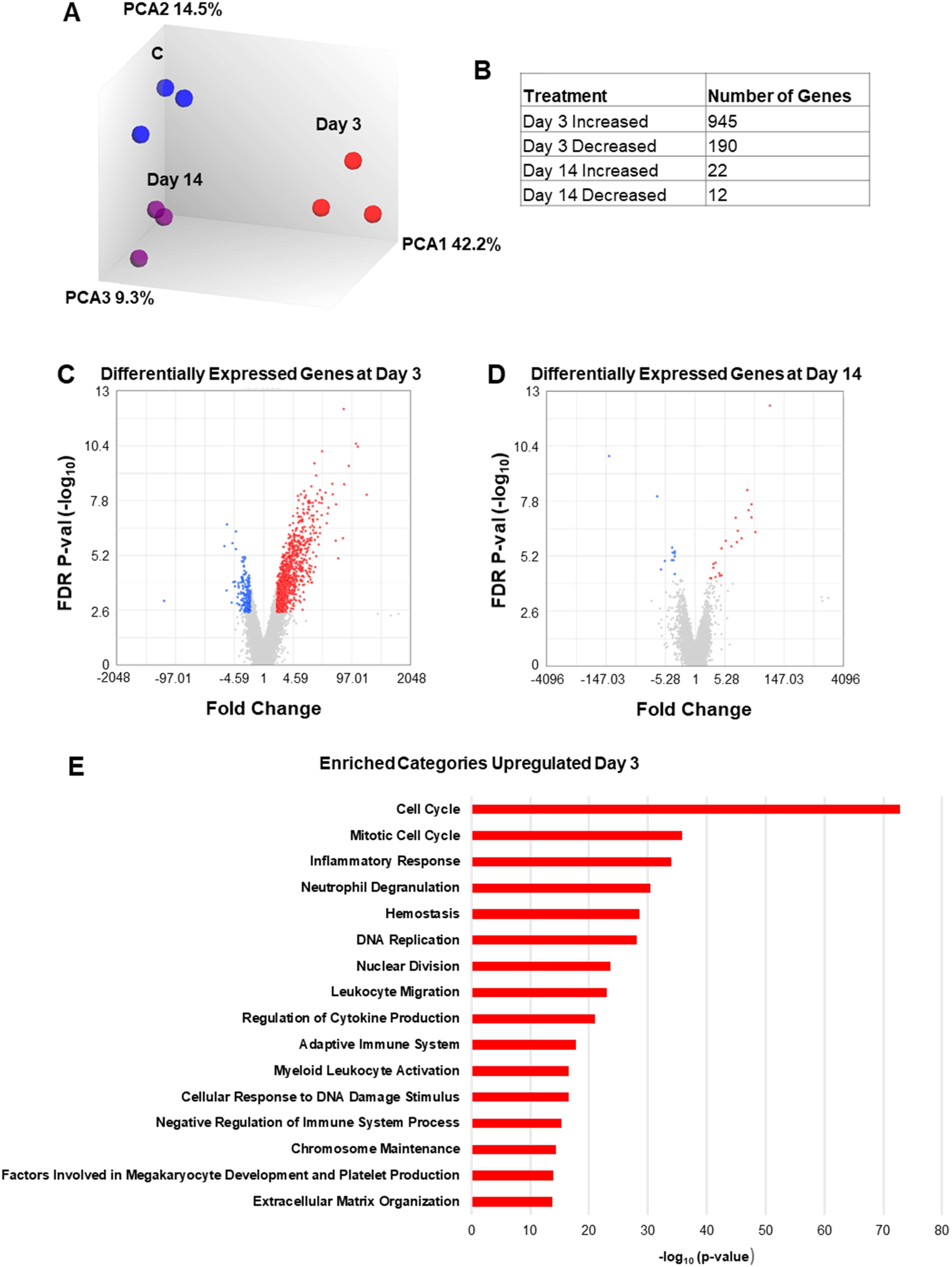
Transcriptome Profiling Reveals Increases in Cell Proliferation, Repair, and Remodeling Processes Three Days After Partial Gland Resection that Resolve After 14 Days. **(A)** Principal component analysis (PCA) transcriptomes from Control (C, blue), Day 3 (red), and Day 14 (purple) shows that each sample is distinct, and triplicates are clustered. **(B)** Total genes changed > 2-fold relative to C were identified using filters for significance (FDR p value<0.05). At Day 3 there many more differentially expressed genes than at Day 14. **(C, D)** Volcano plot generated in Transcriptome Analysis Console (TAC) shows differentially expressed genes at Day 3 (C) and Day 14 (D) relative to C with genes increased greater than 2-fold in red and decreased greater than 2-fold in blue. **(E)** Metascape was used to identify enriched gene categories. The top Reactome (R-MMU) and Gene Ontology (GO) categories are shown. Statistical analysis (in TAC): One-Way Between-Subject p-value and false discovery rate (FDR) p-value using Benjamini-Hochberg procedure. All samples n=3.

### Proliferation is Transiently Upregulated After Partial Gland Resection

As microarray analysis showed that genes involved in cell cycle progression were highly elevated at day 3 but not at day 14 after injury (Appendix Fig. 4), consistent with other organs that undergo an early peak of cell proliferation following partial resection (Voswinckel 2004; Michalopoulos 2010; Kirita et al. 2016), we more closely examined proliferating cells. Since Ki67 can be used to detect cycling cells in any stage of the cell cycle other than G_0_ (Scholzen and Gerdes 2000), we used IHC to detect cells positive for Ki67 (Ki67^+^) in SMGs at 0, 3, 14 and 56 days after resection (Fig. 3A). At day 0, we detected a small number of Ki67^+^ cells scattered throughout the gland. At day 3, significantly increased numbers of Ki67^+^ cells were detected relative to control glands with a small increase still evident at day 14 and nearly returned to control levels by day 56 (Fig. 3A, B). We quantified the cell proliferation rate using Ki67^+^ cells normalized to total nuclei (DAPI^+^). We found that 18 percent of the cells were positive for Ki67 at day 3 after resection (Fig. 3B), which is a 15 fold increase in Ki67^+^ cells relative to control (Appendix Fig. 4). These data indicate that there is a global glandular response to partial gland resection that transiently stimulates cell cycle entry 3 days after injury, but cell proliferation is not sustained.

**Figure 3.**
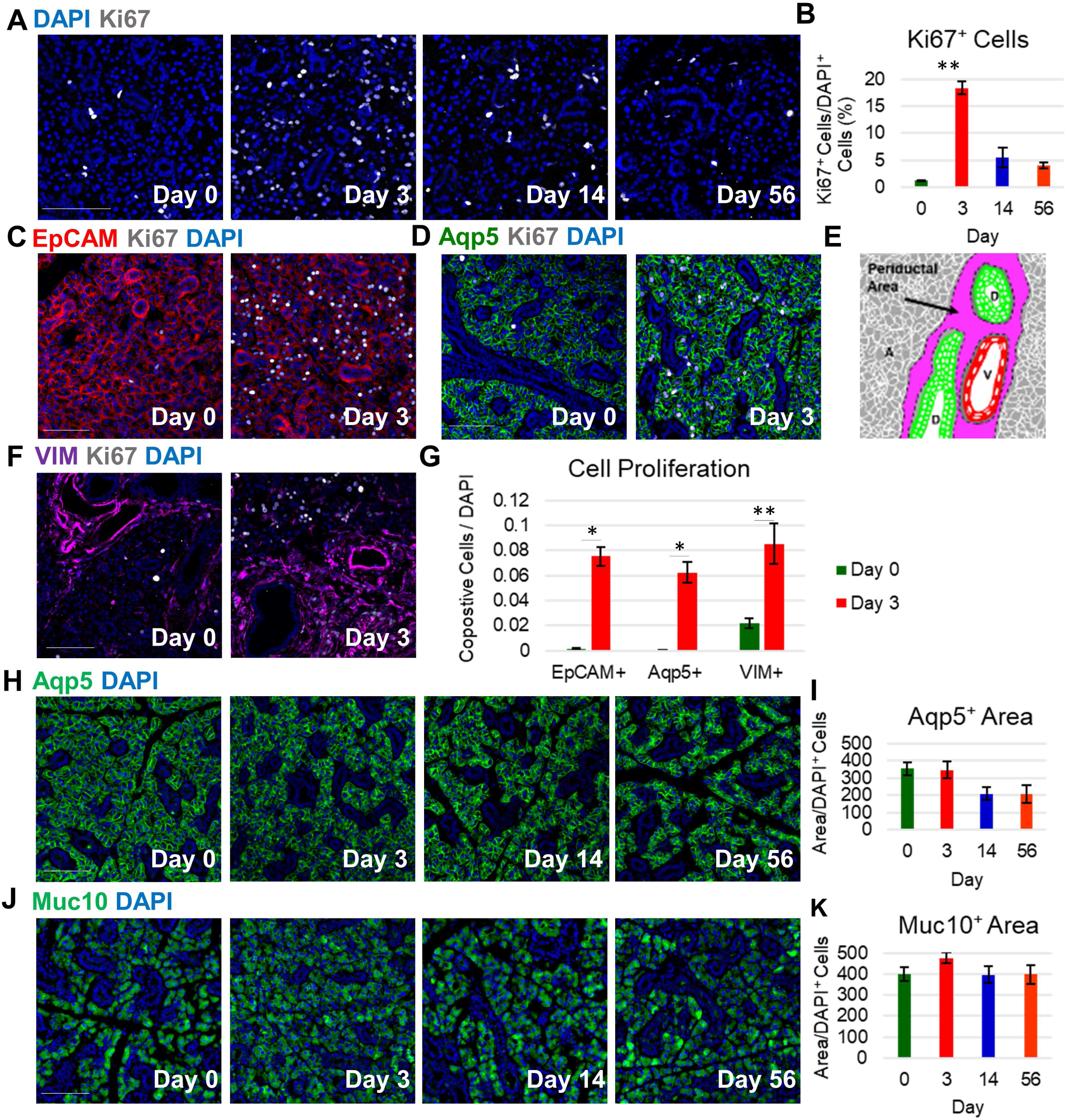
Cell Cycle Entry is Transiently Increased Globally in Both Secretory Epithelial and Stromal Cells, and the Secretory Epithelium is Globally Maintained After Partial Gland Resection. **A)** IHC was performed to detect the proliferation marker, Ki67 (White), relative to DAPI (blue) after partial resection. Ki67^+^ cells are distributed throughout the gland and increase at 3 days after resection. **B)** Quantification of Ki67^+^ cells normalized to total DAPI^+^ cells shows that proliferation significantly increases at 3 days and then declines with 18% of the cells Ki67^+^ 3 days after resection. Staining for Ki67^+^ cells, Day 0 (n=3 glands), Day 3 (n=3), Day 14 (n=4), Day 56 (n=4). Statistical test: One way ANOVA Tukey’s HSD **p≤0.01 **C)** EpCAM (red) and Ki67 (white) IHC at Day 0 and Day3. **D)** Aqp5 (green) and Ki67 (white), Copositive proliferation counts using EpCAM (red) staining with Ki67 (white) IHC at Day 0 and Day3. **E)** Diagram showing periductal area (I, purple), acini (A, gray), ducts (D, green) and vasculature (V, red). **F)** Vimentin (vim, purple) and Ki67 at Day 0 and Day 3. The majority of vim^+^/Ki67^+^ cells are in the periductal regions. **G)** Quantification of the number of Ki67+ cells shows an increase in EpCam^+^, Aqp5^+^ and Vim^+^ cells at Day 3. All copositve cell counts were normalized to DAPI^+^ cells. Images from 3 glands were used for all quantification except for VIM images (n=4). Statistical test: Student’s Two Tailed T-Test, * p≤0.05, ** p≤0.01. **H)** Aqp5 (green) staining in the gland shows that secretory Aqp5^+^ cells are maintained at Day 3.1) Quantification of the area covered by Aqp5 reveals that 35% of the area is lost by Day 14. **J)** IHC for Muc10 (green) shows consistent Muc10 levels across the gland at all time points. **K)** Quantification of the area covered by Muc10 shows no change after resection. IHC for Muc10 and Aqp5 areas, Day 0 (n=4 glands), Day 3 (n=3), Day 14 (n=4), Day 56 (n=4). Error bars are S.E.M. Scale bars, 100 microns.

To examine which cells were responding with a proliferative response we performed IHC to detect Ki67^+^ together with cell type markers at day 3 after resection when Ki67 levels were the highest. We first examined Ki67^+^epithelial cells, performing IHC using an antibody detecting the epithelial cell marker, EpCAM, together with Ki67 (Fig. 3C). The number of Ki67^+^/EpCAM^+^ cells were increased relative to control glands throughout the resected gland (Fig. 3G). To examine the response of secretory acinar cells, we examined Ki67^+^ cells expressing the water channel protein, aquaporin 5 (AQP5) which is expressed in proacinar and acinar cells in salivary epithelium (Nelson et al. 2013) (Fig. 3D). The number of AQP5^+^/Ki67^+^ cells were also increased relative to control glands (Fig. 3G). Indicating that the proliferating epithelial cells were largely secretory acinar cells. To determine if stromal cells, which are enriched in the periductal areas (Fig. 3E) were proliferating, we stained with Ki67 and the mesenchymal marker vimentin (VIM^+^) (Fig. 3F), VIM^+^/Ki67^+^ cells were also increased relative to control glands. Together these data indicate that both the epithelial and stromal cells undergo a global transient proliferative response to injury.

As the secretory capacity of the gland can be reduced by injury, we examined the secretory compartment over time with IHC. IHC for AQP5 and mucin 10 (Muc10) showed persistent expression of these acinar markers through day 56 (Fig. 3H, J). To quantify, we measured the area that was AQP5-or Muc10-positive from thresholded images, and normalized to the number of DAPI^+^ cells. Globally, the AQP5^+^ area decreased by 14 and stabilized by day 56 (Fig. 3I) while Muc10^+^ expressing areas of the gland were sustained (Fig. 3K). Transcriptome analysis to detect SMG secretory genes previously identified from RNA Seq studies (Gao et al. 2018) indicated that mRNAs encoding major secretory products were sustained at day 3 and 14 in resected glands (Appendix Fig. 5). Together, these data suggest that the resected gland maintains functional secretory cells after partial gland resection.

### Secretory Capacity is Lost in Small Local Regions of the Gland

To examine tissue architecture, we performed hematoxylin and eosin (H&E) staining of partially resected glands in comparison with control glands and detected notable regional differences. We observed localized regions with a different morphology (Fig. 4B) than the rest of the gland (Fig. 4A), characterized by increased numbers of small ducts, fewer acini, and more stromal tissue. These small, localized regions were proximal to the cut site and distal to the sublingual gland and were detected in all of the partially resected glands at day 3 and day 14, and 3/4 of the resected glands at day 56 (shown schematically in Fig. 4C). To confirm a reduction in acini in these regions, we measured the tissue area positive for AQP5 and Muc10 in these specific (local) regions (Fig. 4D, F). Loss of both markers occurred at day 3 that became significant and sustained at day 14 through day 56 (Fig. 4E, G). These data indicate reduced functional secretory capacity in these local aberrant regions of the resected glands.

**Figure 4.**
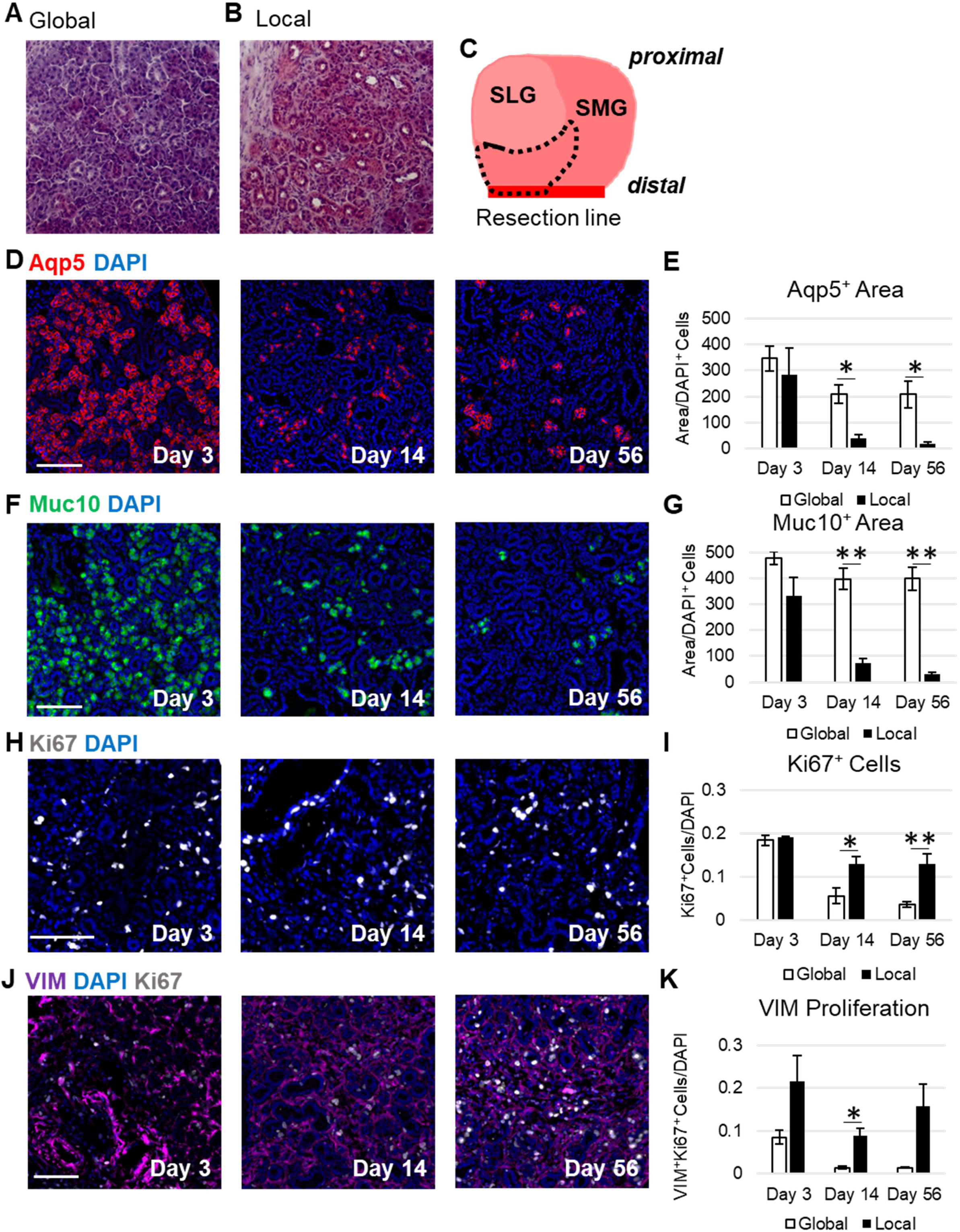
Secretory Capacity is Lost in Local Regions of the Gland and Stromal Proliferation is Increased. **A, B)** H&E Staining shows normal morphology (global) with an aberrant, ductalized pattern in (local) regions of the tissue at Day 14 after resection. **C)** Cartoon depicting the general location of the local tissue response (dotted line) with respect to the sublingual gland (SLG), submandibular gland (SMG), and resection line (red dotted line). **D)** Aqp5^+^ (red) IHC and DAPI (blue) in local aberrant region. **E)** Quantification of Aqp5^+^ area shows a significant decrease within the local region compared to the global. **D)** Muc10^+^cells (green) by IHC and DAPI (blue). Quantification of Muc10^+^ area normalized to DAPI shows a significant loss in the local aberrant region at Day 14 that is not restored by Day 56. **H)** Ki67^+^ cells (white) and DAPI (blue). I) Quantification of Ki67^+^ cells show elevated proliferation in the local region relative to global at both 14 and 56 days after partial gland resection. **J)** Vim (purple) and Ki67 by IHC. **K)** Quantification shows an increase in the number of vim^+^/Ki67^+^ copostive cells at all time points in the local region compared to the global with a significant increase at Day 14. Statistical Analysis: Student’s two-tailed T-Test, *p≤0.05, **p≤0.01. Global measurements Day 3, n=3 glands, day 14, n=4, Day 56 n=4. Local measurements for Ki67: Days 3, 14 (n=4), and Day 56, n=3. Error bars, S.E.M. Scale bars, 100 microns.

### Cell Proliferation is Maintained in Local Regions of the Gland

To investigate cell proliferation specific to these localized regions, we counted the number of Ki67^+^ cells. Interestingly, while global Ki67^+^ cells declined progressively at day 14 and 56 after resection (Fig. 3A, B), we observed a significantly higher number of Ki67^+^ cells maintained at both day 14 and 56 after resection (Fig 4. H, I). With the loss of secretory epithelial markers in the local region, we investigated if these Ki67^+^ cells were mesenchymal. We counted the VIM^+^ and Ki67^+^ (Fig. 4J) copositive cells in the local and global regions. The local Ki67^+^/VIM^+^ cells were maintained at significantly increased numbers compared to the rest of the gland at day 14 (Fig. 4K) and, at all time points VIM^+^ area was higher in the local region than in the global tissue (Appendix Fig. 6), suggesting that the fibroblast injury response is sustained within the specific local region of the gland.

### Regional Differences in Macrophage Abundance and Fibrosis After Partial Gland Resection

Macrophages are important mediators of wound healing and regeneration that persist in chronic wounds. To examine macrophage abundance in the SMG after resection we performed IHC to detect the glycoprotein F4/80 (Fig. 5A), which is widely used as a marker for multiple classes of macrophages (McKnight et al. 1996) We quantified the area covered by F4/80 in resected glands and observed a transient global increase in abundance of F4/80^+^ area at day 3 that returned to baseline by day 14 (Fig. 5B). In the local region, the area covered by F4/80^+^ signal (Fig. 5C) was significantly increased at day 3, and remained elevated at both day 14 and 56 in comparison to the corresponding global tissues (Fig. 5D). These data indicate that macrophages are activated following partial resection but persist only within the local aberrant region.

To examine fibrosis, we used Masson’s Trichrome, a classical histological stain that stains fibrillar collagens and elastins blue revealing regions with high ECM deposition. Globally, we detected a small increase in trichrome staining at day 3 in resected glands relative to control that does not persist or progress (Fig. 5E, F), consistent with early ECM remodeling and a wound healing response. Examination of genes in the ECM remodeling cluster identified by microarray at 3 day, revealed increased expression of genes encoding ECM proteins detected by trichrome stain, including collagen 1 alpha chain 1 *(COL1a1)* and collagen 3 alpha chain 1 (*COL3a1*) (Appendix Fig. 7). These and other ECM proteins that comprise the key structural components of the ECM, called core Matrisome components (Naba et al. 2016), as well as genes whose products are associated with the ECM and involved in ECM remodeling and ECM-based signaling to cells, called Matrisome-associated proteins, were also elevated at day 3, returning to control levels by day 14 (Appendix Fig. 7). In the localized region that retained F4/80 signal, we observed elevated levels of trichrome staining that persisted to day 14 and 56 (Fig. 5G, H and Appendix Fig. 7), indicative of a fibrotic response that appears and persists in the aberrant local region of the gland after resection.

**Figure 5.**
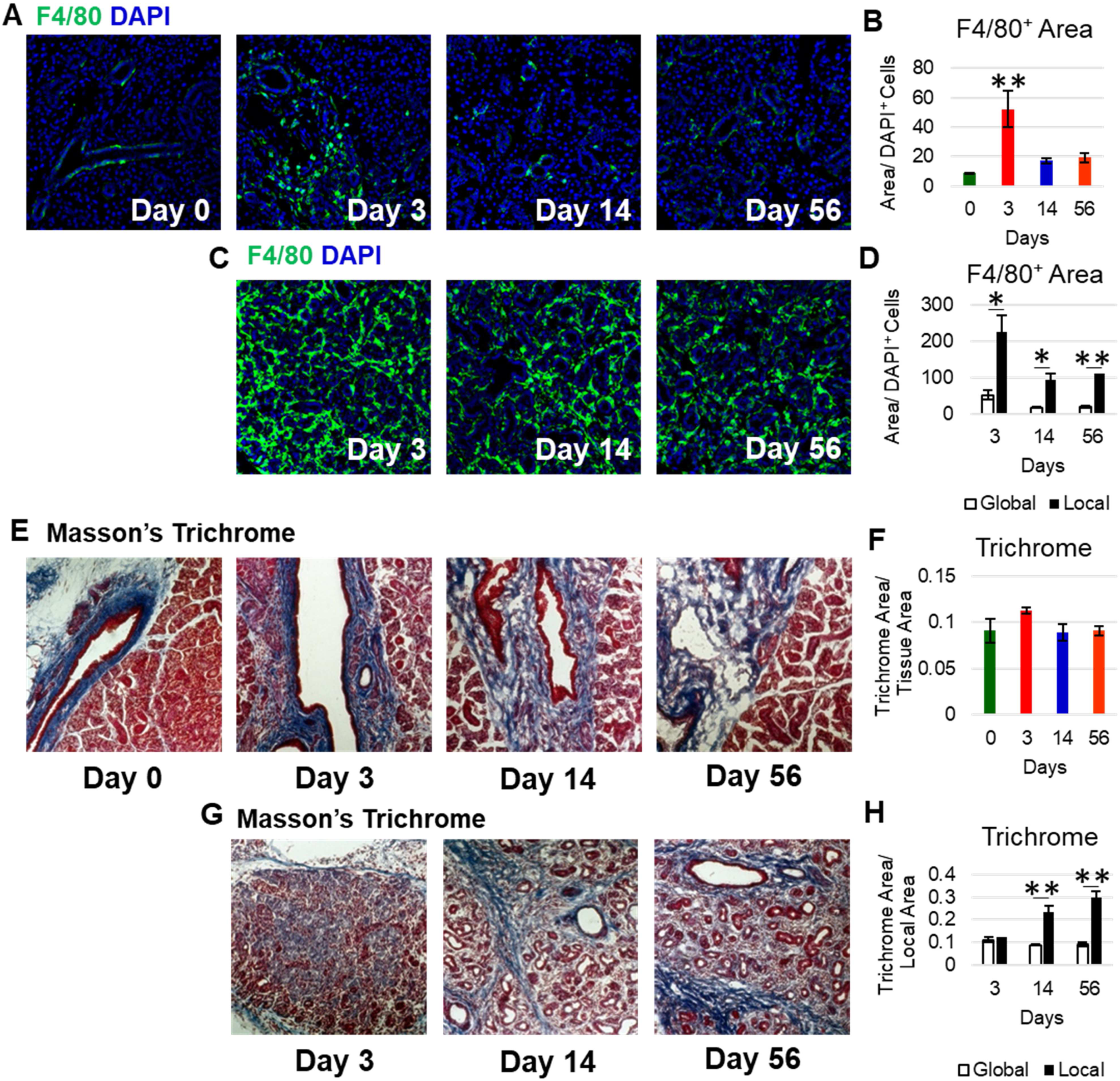
Partial Gland Resection Transiently Increases Macrophages and ECM Deposition Three Days Post Resection Which are Sustained in Localized Aberrant Regions. **A)** IHC for the macrophage marker, F4/80 (green), with DAPI (blue) increased globally in a transient manner at 3 days. **B)** Quantification revealed that the F5/80^+^ area significantly increased 3 days post resection then returned to control levels, as expected with normal wound healing. Statistical Analysis: One way ANOVA Tukey’s HSD **p≤0.01. **C)** F4/80^+^ IHC and DAPI in the local aberrant areas at 3,14, and 56 days. **D)** Quantification revealed a significant increase in F4/80^+^ area in the local aberrant regions relative to the global. **E)** Masson’s Trichrome staining at Days 0, 3,14, and 56. **F)** Quantification of Masson’s Trichrome staining revealed a small, transient increase in ECM in periductal areas 3 days post resection, consistent with an early ECM remodeling response to wound healing. **G)** Masson’s Trichrome stain in local aberrant regions at Days 3, 14, and 56. **H)** Quantification of Masson’s Trichrome stain in the aberrant local regions showed a significant increase in ECM deposition at 14 and 56 days relative to global regions. Statistical Analysis: Student’s Two Tailed T-Test, *p≤0.05, **p≤0.01. Error bars, S.E.M. Scale bars, 100 microns

## Discussion

The SG can partially regenerate after injuries such as ductal ligation and deligation, but does not mount a full regenerative response capable of complete replacement of a functional gland after injury. Using a partial sialoadenectomy model, which lacks other stresses such as the physical pressure induced with ductal ligation and the significant DNA damage induced by irradiation, wherein 40% of the distal tip of the left SMG was removed, we observed a global wound healing response occurring within 3 days that was largely complete day 14 and largely preserved the integrity of the gland. Interestingly, a small localized area of the gland exhibited persistent Ki67 expression, loss of functional secretory acinar cells, increased ECM deposition, and persistent macrophage abundance consistent with a fibrotic response that is reminiscent of more advanced gland pathologies. Regional heterogeneity in salivary gland regenerative capacity has been previously noted by others (van Luijk et al. 2015) but the mechanisms remain poorly understood. This partial gland resection model thus provides the opportunity to study heterogeneous SG repair responses.

Although partial resection of 40% of the mouse SMG does not stimulate a significant regenerative response, the potential of SG cells to regenerate has been demonstrated in studies of ductal ligation and irradiation (Cotroneo et al. 2010; Weng et al. 2018). After deligation, the gland regenerates without intervention, with studies demonstrating that the secretory acinar cells repopulate through a mechanism of clonal expansion by employing lineage tracing of Mist1^+^ acinar progenitor cells (Aure et al. 2015), and involving a process similar to embryonic development (Cotroneo et al. 2010). Regeneration following irradiation can also occur but is typically partial and depends upon the degree of injury (Weng et al. 2018). Multiple mechanisms can limit regenerative ability. Loss of progenitors leads to reduced replacement of lost functional tissue following SG resection and irradiation treatment for head and neck cancer patients (Vissink et al. 2010; Coppes and Stokman 2011), and others. The loss of functional tissue following irradiation, specifically the loss of acinar cells, concomitant with late stage irradiation damage leads to increased infiltrate and fibrosis and further lack of function (Marmary et al., 2016). This partial gland resection model provides a simple model for elucidation of specific cellular mechanisms that limit regeneration. Defining the specific cellular factors that are limiting with each specific type of injury can be applied for development of specific therapeutics.

Cell proliferation is a common response to gland injury with the cell types responding and the degree of response impacting the outcome. Interestingly, the proliferative response in the liver is dependent on the degree of injury, with limited cell proliferation in response to 30% resection versus nearly complete hepatocyte entry into cell cycle after 70% resection (Fausto 2001; Mitchell et al. 2005). After 30% liver resection, around 7% of hepatocytes are BrdU positive while 66% of hepatocytes are after 70% occurring within 3 days that contributes to restoration of liver mass, which is largely complete in both models within 14 days (Miyaoka et al. 2012). We demonstrate that after partial resection of the mouse SMG, cell proliferation occurred in 18% of cells at day 3 but returned to normal levels by day 14, which is similar to the 30% liver resection model that also does not translate to significant increases in tissue mass. Since liver regeneration requires a priming phase to enable efficient response of hepatocytes to growth factor mediated entry into cell cycle and restoration of organ mass, future efforts to improve regeneration of damaged SGs may benefit from identification of priming mechanisms to improve responses to growth factor-mediated stimulation of parenchymal cell proliferation similar to levels observed in successful regeneration paradigms such as ductal ligation studies. Identification of such priming mechanisms could greatly benefit patients with xerostomia resulting from cancer therapy or autoimmune disease, as both cause progressive acinar cell loss that could be remediated by restoration of responsiveness to homeostatic cell regenerative cues. Stimulation of endogenous cells following injury is an appealing therapeutic approach to promote regeneration that avoids complications inherent to cell transplantation therapies, including expansion *in vitro*, transplantation, and engraftment of transplanted cells.

Fibrosis is a component of many clinical conditions that limits organ function. In the salivary gland, fibrosis develops following irradiation treatment for head and neck cancers resulting in loss of functional tissue which is often irreversible (Grundmann et al. 2009; Cheng et al. 2011; Lombaert et al. 2011). Similarly, in the autoimmune disease SjS, fibrosis increases in the salivary glands concomitant with loss of secretory acinar cells as disease progression occurs (Vitali 2002; Konttinen et al. 2006; Bookman et al. 2011; Llamas-Gutierrez et al. 2014; Gervais et al. 2015). Salivary hypofunction in patients with SjS is correlated with an increase in immune infiltrate into the glands and the appearance of fibrosis (Leehan et al. 2018). Interestingly, fibrosis is a hallmark not only of SjS, but also of non-SjS hypofunction, consistent with glandular fibrosis interfering with gland function (Mulholland et al. 2015; Carubbi et al. 2018 Oct 3). Delineating the mechanisms driving regional wound healing differences that occur in the salivary gland in response to partial gland resection may help to identify etiology specific treatment options for xerostomic patients.

Reduction of fibrosis in the connective tissue compartment to restore a permissive regenerative environment facilitates regeneration in many contexts (Bataller and Brenner 2005; Lindquist and Mertens 2013; Rafii et al. 2013; Tanaka and Miyajima 2016; Cordero-Espinoza and Huch 2018). Whether remediation of fibrosis would be sufficient to restore function to damaged and diseased salivary glands is unknown, however limiting fibrosis may be beneficial as part of a combination therapy. Although current mouse models do not adequately recapitulate the fibrotic response in patients, the regional fibrotic response that occurs following partial SMG resection may be useful to elucidate mechanisms of salivary gland fibrosis that limit salivary gland regeneration and identify anti-fibrotic agents that promote functional restoration of fibrotic gland tissue. Additionally, delineating the mechanisms driving regional wound healing differences that occur in the SMG in response to partial gland resection may help to elucidate regional differences in gland responses to SjS.

## Acknowledgements

The authors thank Dr. Paolo Forni for use of his Leica DM 4000 B LED microscope system. The authors are grateful to Drs. Catherine E. Ovitt, Marit H. Aure, Thomas Begley, Antigone McKenna, DVM and Sara Evke for helpful suggestions, insight, and assistance in implementing this surgical model. This research was funded by NIH grants R56DE02246706, R21DE027571, and R01DE02795301 (to ML) and funds from the University at Albany, SUNY. KD was partially supported by NIH NRSA fellowship, F32DE027868.

## Author Contributions

K.O., K.D., D.N, and M.L. designed experiments, analyzed data, assembled figures, and wrote and revised the manuscript. K.O., K.D., E.T., and A.A. performed the experiments and analyzed data. A.S. and M.P. performed experiments.

## Conflicts of Interest

The authors declare no financial or non-financial conflicts of interest.

## Data Availability

All data generated or analyzed during this study are included in this published article and its supplementary information files. Microarray data is available at GEO (https://www.ncbi.nlm.nih.gov/geo). Supplementary information is available in Appendix.

## Notes

#### Summary of Updates

higher resoluation images are included and an updated appendix file

https://ncbi.nlm.nih.gov/geo

## References

Aure MH, Konieczny SF, Ovitt CE. 2015. Salivary Gland Homeostasis Is Maintained through Acinar Cell Self-Duplication. Dev Cell. 33(2):231–237. doi:10.1016/j.devcel.2015.02.013.

Bataller R, Brenner DA. 2005. Liver fibrosis. J Clin Invest. 115(2):209–218. doi:10.1172/JCI200524282.

Bookman AAM, Shen H, Cook RJ, Bailey D, McComb RJ, Rutka JA, Slomovic AR, Caffery B. 2011. Whole stimulated salivary flow: Correlation with the pathology of inflammation and damage in minor salivary gland biopsy specimens from patients with primary Sjögren’s syndrome but not patients with sicca. Arthritis Rheum. 63(7):2014–2020. doi:10.1002/art.30295.

Carubbi F, Alunno A, Gerli R, Giacomelli R. 2018 Oct 3. Histopathology of salivary glands. Reumatismo.:146–154. doi:10.4081/reumatismo.2018.1053.

Cheng SCH, Wu VWC, Kwong DLW, Ying MTC. 2011. Assessment of post-radiotherapy salivary glands. Br J Radiol. 84(1001):393–402. doi:10.1259/bjr/66754762.

Coppes R, Stokman M. 2011. Stem cells and the repair of radiation-induced salivary gland damage: Stem cells to repair irradiated salivary glands. Oral Dis. 17(2):143–153. doi:10.1111/j.1601-0825.2010.01723.x.

Cordero-Espinoza L, Huch M. 2018. The balancing act of the liver: tissue regeneration versus fibrosis. J Clin Invest. 128(1):85–96. doi:10.1172/JCI93562.

Cotroneo E, Proctor GB, Carpenter GH. 2010. Regeneration of acinar cells following ligation of rat submandibular gland retraces the embryonic-perinatal pathway of cytodifferentiation. Differentiation. 79(2):120–130. doi:10.1016/j.diff.2009.11.005.

Cowan MJ, Crystal RG. 1975. Lung growth after unilateral pneumonectomy: quantitation of collagen synthesis and content. Am Rev Respir Dis. 111(3):267–277. doi:10.1164/arrd.1975.111.3.267.

Fausto N. 2001. Liver regeneration: From laboratory to clinic. Liver Transpl. 7(10):835–844. doi:10.1053/jlts.2001.27865.

Gao X, Oei MS, Ovitt CE, Sincan M, Melvin JE. 2018. Transcriptional profiling reveals gland-specific differential expression in the three major salivary glands of the adult mouse. Physiol Genomics. 50(4):263–271. doi:10.1152/physiolgenomics.00124.2017.

Ge N, Peng X, Zhang L, Cai Z-G, Guo C-B, Yu G-Y. 2016. Partial sialoadenectomy for the treatment of benign tumours in the submandibular gland. Int J Oral Maxillofac Surg. 45(6):750–755. doi:10.1016/j.ijom.2015.12.013.

Gervais EM, Desantis KA, Pagendarm N, Nelson DA, Enger T, Skarstein K, Liaaen Jensen J, Larsen M. 2015. Changes in the Submandibular Salivary Gland Epithelial Cell Subpopulations During Progression of Sjögren’s Syndrome-Like Disease in the NOD/ShiLtJ Mouse Model: Epithelial Subpopulations in NOD/ShiLtJ Mouse. Anat Rec. 298(9):1622–1634. doi:10.1002/ar.23190.

Grundmann O, Mitchell GC, Limesand KH. 2009. Sensitivity of Salivary Glands to Radiation: from Animal Models to Therapies. J Dent Res. 88(10):894–903. doi:10.1177/0022034509343143.

Kirita Y, Kami D, Ishida R, Adachi T, Tamagaki K, Matoba S, Kusaba T, Gojo S. 2016. Preserved Nephrogenesis Following Partial Nephrectomy in Early Neonates. Sci Rep. 6(1). doi:10.1038/srep26792. [accessed 2019 Jul 23]. http://www.nature.com/articles/srep26792.

Konttinen YT, Porola P, Konttinen L, Laine M, Poduval P. 2006. Immunohistopathology of Sjögren’s syndrome. Autoimmun Rev. 6(1):16–20. doi:10.1016/j.autrev.2006.03.003.

Leehan KM, Pezant NP, Rasmussen A, Grundahl K, Moore JS, Radfar L, Lewis DM, Stone DU, Lessard CJ, Rhodus NL, et al. 2018. Minor salivary gland fibrosis in Sjögren’s syndrome is elevated, associated with focus score and not solely a consequence of aging. Clin Exp Rheumatol. 36 Suppl 112(3):80–88.

Lindquist JA, Mertens PR. 2013. Myofibroblasts, regeneration or renal fibrosis--is there a decisive hint? Nephrol Dial Transplant. 28(11):2678–2681. doi:10.1093/ndt/gft247.

Llamas-Gutierrez FJ, Reyes E, Martínez B, Hernández-Molina G. 2014. Histopathological environment besides the focus score in Sjögren’s syndrome. Int J Rheum Dis. 17(8):898–903. doi:10.1111/1756-185X.12502.

Lombaert I, Knox S, Hoffman M. 2011. Salivary gland progenitor cell biology provides a rationale for therapeutic salivary gland regeneration: Salivary gland regeneration using progenitor cells. Oral Dis. 17(5):445–449. doi:10.1111/j.1601-0825.2010.01783.x.

Lombaert I, Movahednia MM, Adine C, Ferreira JN. 2017. Concise Review: Salivary Gland Regeneration: Therapeutic Approaches from Stem Cells to Tissue Organoids: Translational and Clinical Research. STEM CELLS. 35(1):97–105. doi:10.1002/stem.2455.

van Luijk P, Pringle S, Deasy JO, Moiseenko VV, Faber H, Hovan A, Baanstra M, van der Laan HP, Kierkels RGJ, van der Schaaf A, et al. 2015. Sparing the region of the salivary gland containing stem cells preserves saliva production after radiotherapy for head and neck cancer. Sci Transl Med. 7(305):305ra147–305ra147. doi:10.1126/scitranslmed.aac4441.

Marmary Y, Adar R, Gaska S, Wygoda A, Maly A, Cohen J, Eliashar R, Mizrachi L, Orfaig-Geva C, Baum BJ, et al. 2016. Radiation-Induced Loss of Salivary Gland Function Is Driven by Cellular Senescence and Prevented by IL6 Modulation. Cancer Res. 76(5):1170–1180. doi:10.1158/0008-5472.CAN-15-1671.

McKnight AJ, Macfarlane AJ, Dri P, Turley L, Willis AC, Gordon S. 1996. Molecular Cloning of F4/80, a Murine Macrophage-restricted Cell Surface Glycoprotein with Homology to the G-protein-linked Transmembrane 7 Hormone Receptor Family. J Biol Chem. 271(1):486–489. doi:10.1074/jbc.271.1.486.

Michalopoulos GK. 2010. Liver Regeneration after Partial Hepatectomy. Am J Pathol. 176(1):2–13. doi:10.2353/ajpath.2010.090675.

Mitchell C, Nivison M, Jackson LF, Fox R, Lee DC, Campbell JS, Fausto N. 2005. Heparin-binding Epidermal Growth Factor-like Growth Factor Links Hepatocyte Priming with Cell Cycle Progression during Liver Regeneration. J Biol Chem. 280(4):2562–2568. doi:10.1074/jbc.M412372200.

Miyaoka Y, Ebato K, Kato H, Arakawa S, Shimizu S, Miyajima A. 2012. Hypertrophy and Unconventional Cell Division of Hepatocytes Underlie Liver Regeneration. Curr Biol. 22(13):1166–1175. doi:10.1016/j.cub.2012.05.016.

Mulholland GB, Jeffery CC, Satija P, Côté DWJ. 2015. Immunoglobulin G4-related diseases in the head and neck: a systematic review. J Otolaryngol - Head Neck Surg. 44(1). doi:10.1186/s40463-015-0071-9. [accessed 2019 Jul 24]. https://journalotohns.biomedcentral.com/articles/10.1186/s40463-015-0071-9.

Naba A, Clauser KR, Ding H, Whittaker CA, Carr SA, Hynes RO. 2016. The extracellular matrix: Tools and insights for the “omics” era. Matrix Biol. 49:10–24. doi:10.1016/j.matbio.2015.06.003.

Nelson DA, Manhardt C, Kamath V, Sui Y, Santamaria-Pang A, Can A, Bello M, Corwin A, Dinn SR, Lazare M, et al. 2013. Quantitative single cell analysis of cell population dynamics during submandibular salivary gland development and differentiation. Biol Open. 2(5):439–447. doi:10.1242/bio.20134309.

Nevzorova Y, Tolba R, Trautwein C, Liedtke C. 2015. Partial hepatectomy in mice. Lab Anim. 49(1_suppl):81–88. doi:10.1177/0023677215572000.

Rafii R, Juarez MM, Albertson TE, Chan AL. 2013. A review of current and novel therapies for idiopathic pulmonary fibrosis. J Thorac Dis. 5(1):48–73. doi:10.3978/j.issn.2072-1439.2012.12.07.

Scholzen T, Gerdes J. 2000. The Ki-67 protein: From the known and the unknown. J Cell Physiol. 182(3):311–322. doi:10.1002/(SICI)1097-4652(200003)182:3<311::AID-JCP1>3.0.CO;2-9.

Tanaka M, Miyajima A. 2016. Liver regeneration and fibrosis after inflammation. Inflamm Regen. 36(1). doi:10.1186/s41232-016-0025-2. [accessed 2019 Jul 23]. http://inflammregen.biomedcentral.com/articles/10.1186/s41232-016-0025-2.

Vissink A, Mitchell JB, Baum BJ, Limesand KH, Jensen SB, Fox PC, Elting LS, Langendijk JA, Coppes RP, Reyland ME. 2010. Clinical Management of Salivary Gland Hypofunction and Xerostomia in Head-and-Neck Cancer Patients: Successes and Barriers. Int J Radiat Oncol. 78(4):983–991. doi:10.1016/j.ijrobp.2010.06.052.

Vitali C. 2002. Classification criteria for Sjogren’s syndrome: a revised version of the European criteria proposed by the American-European Consensus Group. Ann Rheum Dis. 61(6):554–558. doi:10.1136/ard.61.6.554.

Voswinckel R. 2004. Characterisation of post-pneumonectomy lung growth in adult mice. Eur Respir J. 24(4):524–532. doi:10.1183/09031936.04.10004904.

Weng P-L, Aure MH, Maruyama T, Ovitt CE. 2018. Limited Regeneration of Adult Salivary Glands after Severe Injury Involves Cellular Plasticity. Cell Rep. 24(6):1464–1470.e3. doi:10.1016/j.celrep.2018.07.016.

Xiong J, Hou J. 2016. Apical Resection Mouse Model to Study Early Mammalian Heart Regeneration. J Vis Exp.(107). doi:10.3791/53488. [accessed 2019 Jul 23]. http://www.jove.com/video/53488/apical-resection-mouse-model-to-study-early-mammalian-heart.

Zhou Y, Zhou B, Pache L, Chang M, Khodabakhshi AH, Tanaseichuk O, Benner C, Chanda SK. 2019. Metascape provides a biologist-oriented resource for the analysis of systems-level datasets. Nat Commun. 10(1). doi:10.1038/s41467-019-09234-6. [accessed 2019 Jul 23]. http://www.nature.com/articles/s41467-019-09234-6.

