## Appendix for "Regional Differences Following Partial Salivary Gland Resection"

**Appendix Materials and Methods**

***Partial Sialoadenectomy***

Female C57Bl6/J mice were housed in accordance with husbandry protocols at the University at Albany in a 12 hour light/dark cycle until reaching 12-16 weeks of age, mice were progeny of mice obtained from The Jackson Laboratory. Female mice were chosen due to similarities in tissue architecture with human salivary glands (Amano et al. 2012; Maruyama et al. 2019). All animal procedures were performed in accordance with protocols approved by the University at Albany, SUNY IACUC committee. At 12 weeks of age, we performed unilateral surgical resection of approximately 40% the left submandibular salivary gland under 2% isoflurane anesthesia. Surgeries were performed mid-day. Mice were continuously monitored under anesthesia, monitored for pain, distress, and weight loss or gain post-surgery. Mice for each surgery were chosen at random prior to surgery and housed singly until day 14 then housed in groups for longer time points. We immediately harvested the gland, day zero, or allowed a recovery period of 3, 14 or 56 days before harvesting glands. Mice were euthanized under CO_2_ in accordance with euthanasia procedures at the University at Albany. We also quantified glands of age- and gender-matched control mice. Buprenorphine (0.015mg/mL) was administered post-operatively, subcutaneously as an analgesic. Incisions were closed with three to five interrupted sutures. Gland dimensions were measured in the animals post-euthanasia with calipers prior to gland collection, and weights were measured with the sublingual gland (SLG) attached. Mice were assigned a unique identifier at the time of surgery that was used to identify the surgical procedure and treatment. All subsequent sample processing, image capture, and quantitative analysis was performed blinded with reference only to the unique identifiers.

***Preparation of Cryosections***

Glands were removed and fixed in 4% paraformaldehyde in 1X phosphate buffered saline (4% PFA-PBS) for 2 hours, washed, incubated in an increasing sucrose gradient and left in 30% sucrose overnight at 4C. Glands were then transferred to 15% sucrose in optimal cutting temperature (OCT) overnight at 4C, then frozen over liquid nitrogen. Ten micron cryosections were serially sectioned and dried on SuperFrost Plus glass slides for 30 minutes (Electron Microscopy Sciences). Slides were stored at -80C. After removal from -80C, slides were gradually allowed to return to room temperature.

***Fluorescent Immunohistochemistry***

Slides were post-fixed in 4% PFA-PBS for 18 minutes, followed by rinsing in PBS. Slides were permeabilized in 0.5% Triton X-100 in PBS for 18 minutes, followed by two rinses in PBS. Blocking was performed in 3% bovine serum albumin 0.5% Triton X-100 in PBS (3% BSA-0.5% Tx-PBS) for 90 minutes at room temperature in a humidified chamber. Antibodies were diluted in 3% BSA-PBS and applied for a minimum of one hour at room temperature or overnight at 4C (Appendix Table 1). Four PBS washes were performed after application of each round of primary antibody. Secondary antibodies or direct conjugates (Appendix Table 1) were added in 3% BSA-0.5% Tx-PBS for 90 minutes, after application two PBS washes were performed, followed by DAPI staining for 10 min, and two additional PBS washes. Multiplexed immunohistochemistry was performed as described previously(Nelson et al. 2013), with the exception that 0.1M sodium citrate pH6.0 was used for antigen retrieval. Coverslips were applied with a glycerol-based mounting media containing 4% n-propyl galate as an antifade, as previously reported(Nelson et al. 2013).

***Hematoxylin and Eosin Staining***

Room temperature cryosections were incubated in PBS for 5 minutes. Slides were incubated in Modified Harris Hematoxylin (Fisher Scientific) for 1 minute and rinsed in running tap water for 1 minute. Slides were then incubated in Eosin Y, 1% aqueous, (Electron Microscopy Sciences) for 1 minute and rinsed in running tap water for one minute. Slides were then dehydrated in 100% ethanol for 3 minutes and cleared in xylene for 30 seconds. Slides were then mounted with permount (Fisher Scientific).

***Masson’s Trichrome Staining***

Room temperature cryosections were fixed in Bouin’s Fixative (Polysciences) for 20 minutes. Slides were rinsed in cold tap water for 5 minutes, then stained in Weigert’s Iron Hematoxylin (Polysciences) for 5 seconds. Slides were rinsed in cold tap water for 2 minutes, then stained in Beibrich Scarlett-Acid Fuchsin Solution (Polysciences) for 15 seconds. Slides were rinsed in deionized water (DI) until the water running off of the slide was clear. Phosphotungstic/Phosphomolybdic acid (Polysciences) was added directly to the tissue sections and incubated for 10 minutes then drained onto a Kimwipe. Slides were stained in Aneline Blue (Polysciences) for 30 seconds and rinsed in DI three times for 30 second each. 1% Acetic Acid (Polysciences) was pipetted directly onto the seconds for 1 minute then rinsed in DI. Slides were dehydrated in 95% and 100% ethanol for 2 minutes each and cleared in xylene for 2 minutes then mounted in Permount (Fisher Scientific).

***Image Processing, Quantification, and Statistical Analysis***

For quantification of each marker by IHC or histological stain, sections obtained from similar tissue depths and similar regions of the gland were compared. Sections having low tissue quality due to poor fixation or tissue loss were excluded from analysis. IHC sections were imaged on an Olympus IX-81 microscope with a Peltier-cooled CCD camera (Q imaging, RET 4000DC-F-M-12) using a PLAN S-APO 20X, 0.75 NA objective. Sections stained with histological stains were imaged on a Leica DM 4000 B LED scope with a Leica DFC 310 FX RGB color camera or on a Nikon TS100 inverted microscope fitted with a Canon. All image processing and quantification was performed using the freeware FIJI (Schindelin et al. 2012).

For trichrome analysis, control and resected tissues were examined as whole tissues with n=3 or 4 for each group. For quantification of Masson’s Trichrome, areas staining blue were selected with color thresholding performed equally across groups and quantified. Color thresholding was used to quantify total tissue area per sample. Positive staining area was normalized to total tissue area.

Estimated DAPI^+^ nuclei were calculated by autothresholding DAPI and performing particle analysis using Fiji. 10 single DAPI^+^ particles were used to create a minimum and maximum single DAPI profile. Particles were then counted if they fell within the profile for single DAPI and divided by the average DAPI area if they were larger than the maximum area for a single nucleus, signifying the presence of multiple nuclei within a particle. Numbers of cells within images were averaged per tissue piece (Appendix Figure 2). To quantify the number of Ki67^+^ cells, cells were automatically counted using the same settings as the estimated DAPI^+^ nuclei. To measure differences in the local areas, areas around the local tissue were looped out, cleared, and inverted in Fiji to only measure the local area. Local areas were determined by their abnormal tissue structure. (Fig. 4B, C). Aqp5^+^ and Muc10^+^ areas were quantified using equally thresholded images and whole images were measured. Areas were then normalized to estimated DAPI^+^ cells.

Aqp5^+^/Ki67^+^ cells and Epcam^+^/Ki67^+^ cells total copositive cells were counted on 20x images. Copositive cells were counted n=3 in control and 3 days post resection and normalized to the estimated number of DAPI^+^ cells. VIM^+^/Ki67^+^ total copositive cells were counted in 5 randomly generated ROIs from 3 images closest to the mean VIM^+^ area. VIM^+^/Ki67^+^ copositive cells in the localized region were counted by counting randomly generated ROIs that were created in the localized region. All VIM^+^/Ki67^+^ copositive cell counts were normalized to the estimated number of DAPI^+^ cells within the ROIs.

All tabulation of quantification was performed in Microsoft Excel. Statistical analysis was performed using either Microsoft Excel or freeware VassarStats (Lowry 2008). Either a two-tailed Students t-test or an ANOVA followed by Tukey’s multiple comparisons test was performed indicated specifically in figure legends. A sample size of 3 animals were used for each treatment as statistical relevance was achieved with this n, reducing the number of animals needed for the study, when appropriate, larger sample sizes were used.

***RNA Isolation***

Total RNA was isolated from day 3, day 14, and mock surgery control mice using TRIzol (Ambion) /chloroform (Fisher Scientific). All centrifuge steps were completed at 16,000 x g using the Eppendorf 5415C centrifuge. Manufacturer’s protocol was followed for sample lysis with one additional chloroform step added after isolation of the aqueous layer. RNA was precipitated using 2 volumes of 100% molecular grade ethanol (Sigma) and 1/10 volume of 3M sodium acetate (pH=5.2) and incubated for 2 hours at -80C. The manufacturer’s protocol for washing RNA was repeated twice prior to air drying the pellet. Manufacturer’s protocol was used for solubilizing the RNA. Concentrations were determined using a NanoDrop ND-1000 Spectrophotometer. To remove genomic DNA, extracted total RNA was treated with the Turbo DNA-free^TM^ Kit (Invitrogen), following the manufacturer’s protocol for routine DNase treatment procedure in a 30 μL total volume. RNeasy Plus Mini Kit (Qiagen) was then used to further purify the RNA following manufacturer’s protocol. RNA quality was evaluated with the Agilent BioAnalyzer all samples had a RIN >7.

***Microarray***

Samples were hybridized to the Clariom S Assay, mouse (Applied Biosystems) at the Center for Functional Genomics, University at Albany, SUNY N=3 for Day 3, Day 14, and mock surgery control. Total RNA (100 ng) was processed using the WT Plus Reagent kit (Affymetrix, Santa Clara, CA). Sense target cDNAs were generated using the standard Affymetrix WT protocol and hybridized to Affymetrix Mouse Clariom S arrays. Arrays were washed, stained and scanned on a GeneChip 3000 7G scanner using Affymetrix GeneChip Command Console Software (AGCC). Transcriptome Analysis Console Software (TAC v4.0.1.36) was used to identify differentially expressed genes. Briefly the CEL files were summarized using the SST-RMA algorithm in TAC and the normalized data were subjected to one-way ANOVA with a Benjamini Hochberg False Discovery Rate correction included (p<0.05). Gene enrichment analysis was performed using Metascape online software (metascape.org; Zhou et al. 2019). Genes with a fold change ≥2 or ≤-2 and an FDR p value<0.05 were input to Metascape Express Analysis.

**Appendix Figure 1. Contralateral Glands Respond to Partial Resection by Changing Length, Width, and Weight. A)** Lengths of contralateral glands (mm) measured *in situ* with calipers, demonstrate that glands respond to resection of the left gland by increasing in length up through day 56. **B)** Widths of contralateral glands (mm) measured *in situ* with calipers, show that glands increase in width, reaching a maximum 14 days after resection. **C)** Weights of contralateral glands (mg) demonstrate that glands reach a maximum at Day 14. Control (C) (n=5), resection with immediate tissue harvest (Day 0) (n=9), 3 days post resection (Day 3) (n=11), 14 days post resection (Day 14) (n=8) and 56 days post resection (Day 56) (n=4). Error bars are S.E.M. Statistical Tests: One-Way ANOVA, * p≤0.05, ** p≤0.01

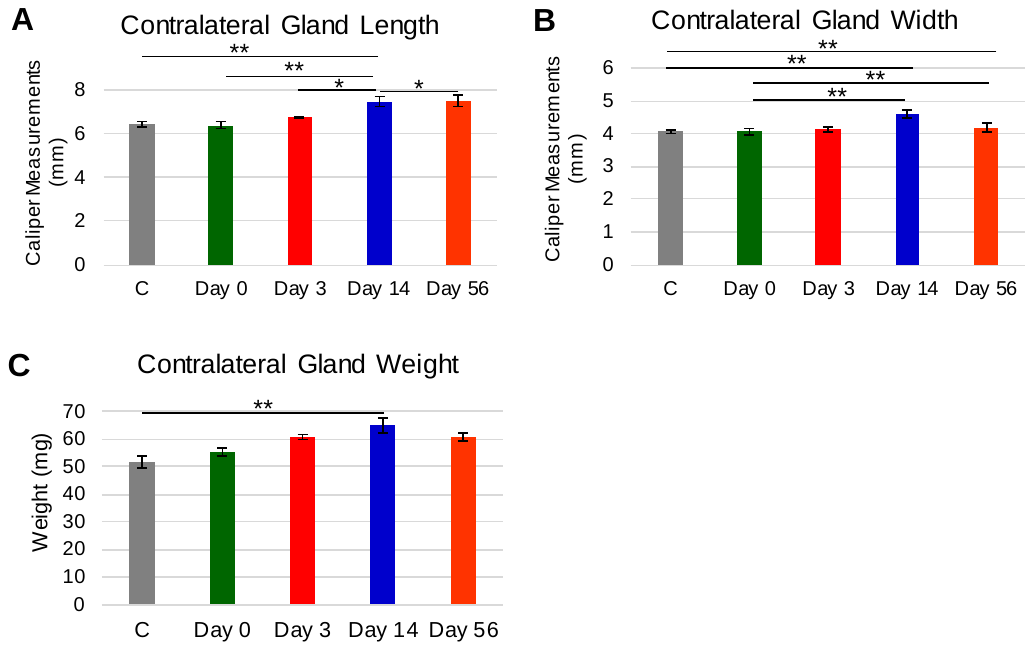

**Appendix Figure 2. Automatic DAPI Cell Counts to Estimate DAPI^+^ Cells. A)** Manual counting of DAPI-positive nuclei in Fiji. Yellow box shows ROI with cyan dots denoting counters. **B)** Automatic thresholding of DAPI-positive nuclei. **C)** Mask created by particle analysis function in Fiji counting DAPI-positive nuclei. **D)** Single DAPI positive particles counted in Excel using the averaged single DAPI nuclei size from ten single nuclei. Multiple DAPI calculated by dividing particles that were larger than one single DAPI by the average area of a single DAPI particle. EST DAPI is the sum of the single DAPI and multiple DAPI. Comparison of EST DAPI and Manual DAPI counting shows that EST DAPI is 94% accurate in estimation of DAPI-positive cells within ROIs.

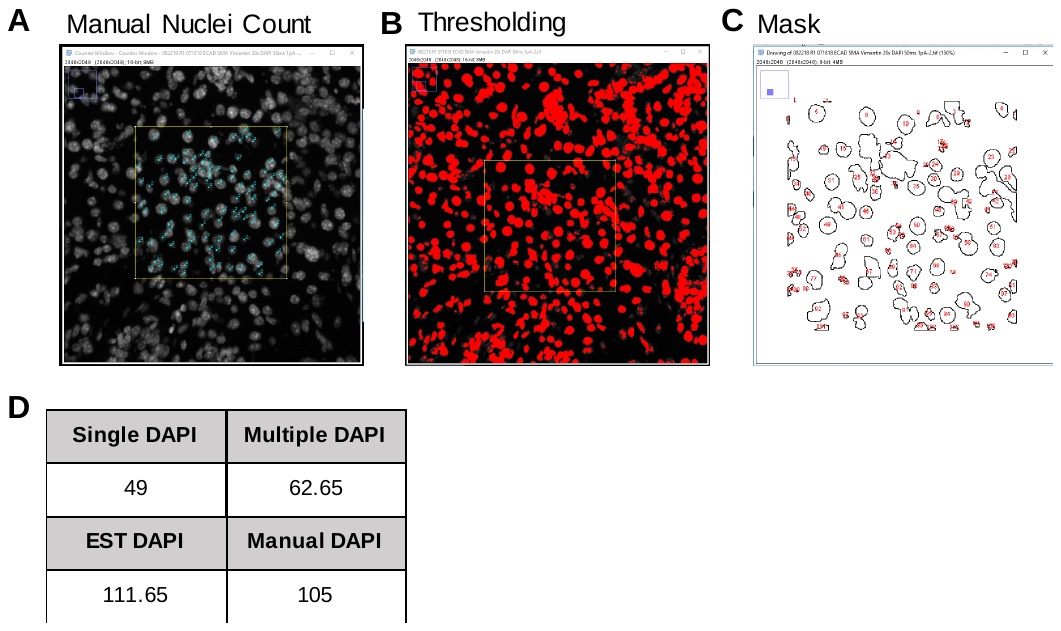

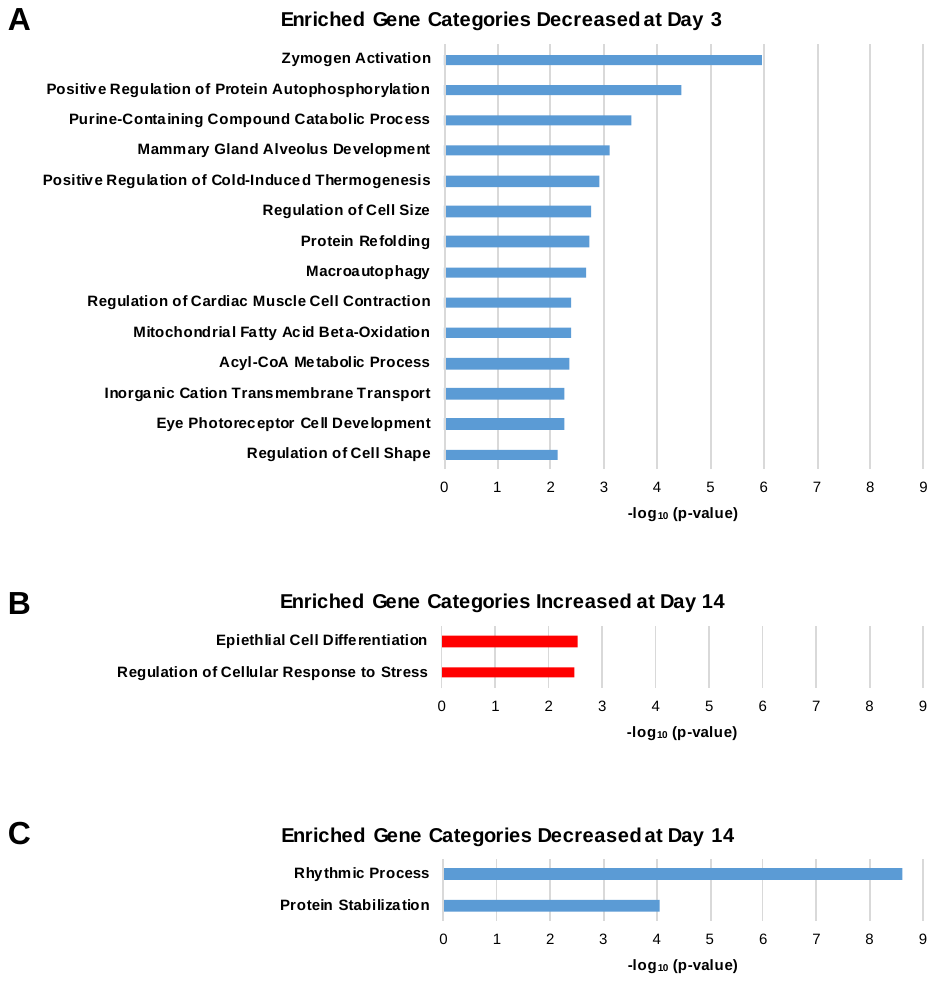

**Appendix Figure 3. Enriched Gene Categories Decreased at Day 3 and Changed at Day 14. A)** Significantly decreased gene categories at Day 3. **B)** Gene categories significantly increased at Day 14 and **C)** Gene categories significantly decreased at Day 14. Both Reactome (R-MMU) and Gene Ontology (GO) categories are shown for all lists. All lists were generated by selecting for genes decreased by 2-fold or greater with FDR, p < 0.05. Enriched categories decreased at Day 3 include genes involved in cellular maintenance and metabolic processes. Gene subsets changed at 3 days are not sustained at 14 days. n=3 biological replicates per condition.

**Appendix Figure 4. Cell Cycle Genes and Ki67 Protein Levels are Increased 3 Days Post Resection. A)** Genes included in the enriched categories Cell cycle (R-MMU-1640170) and Mitotic Cell Cycle (GO:0000278) were used to generate a short gene list of changed cell cycle genes. The top 10 significantly differentially expressed genes at Day 3 were used to generate the table. Changes in cell cycle genes are not maintained at 14 days. **B)** Quantification of Ki67-positive cells shows a 15 fold increase in Ki67 normalized to DAPI at day 3 after resection throughout the entire tissue when normalized to day 0. Levels of Ki67-positive cells taper off at day 14 and 56. Day 0 (n=3), Day 3 (n=3), Day 14 (n=4), Day 56 (n=4). Error bars are S.E.M. Statistical test: One-way ANOVA with Tukey’s HSD post-hoc test. ** p≤ 0.01.

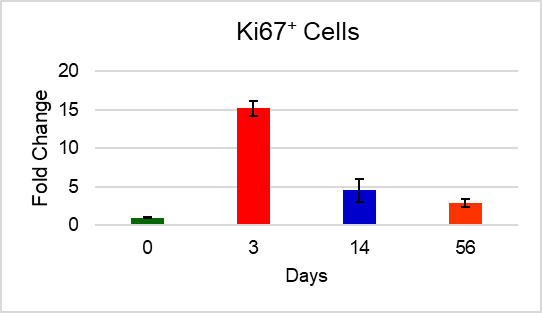

**B**

**

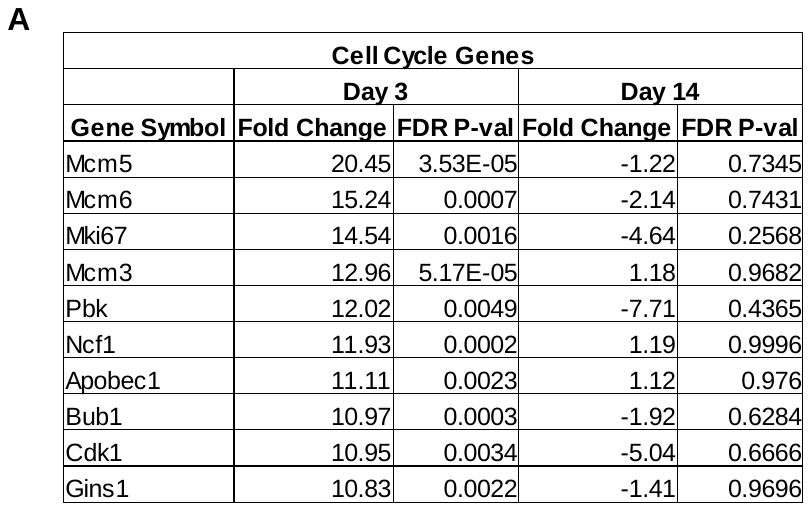

Ki67+ Cells

Fold Change

| Days | Ki67/DAPI |
| --- | --- |
| 0 | 1 |
| 3 | 15.2275 |
| 14 | 4.57629 |
| 56 | 2.92221 |

**Appendix Figure 5. SMG Secretory Genes are not Significantly Changed After Partial Gland Resection.** SMG secretory gene list based on (Gao et al. 2018) at days 3 and 14 reveals little difference in any gene relative to control at either time point.

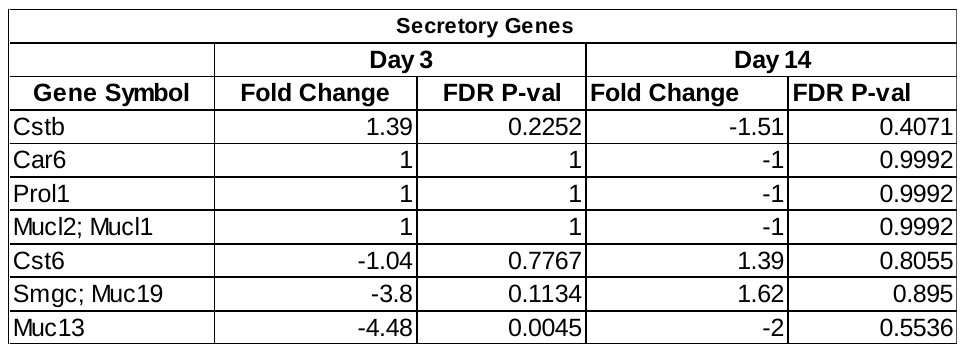

**Appendix Figure 6. VIM^+^ Area Increases at Day 3 and Remains Highly Expressed at Later Time Points.** Quantification of IHC for VIM revealed a significant increase in VIM-positive area in the local aberrant regions relative to the global appearing at 3 days and persisting at days 14 and 56. Statistical Analysis: Students Two Tailed T-Test **p≤0.01. Error bars, S.E.M.

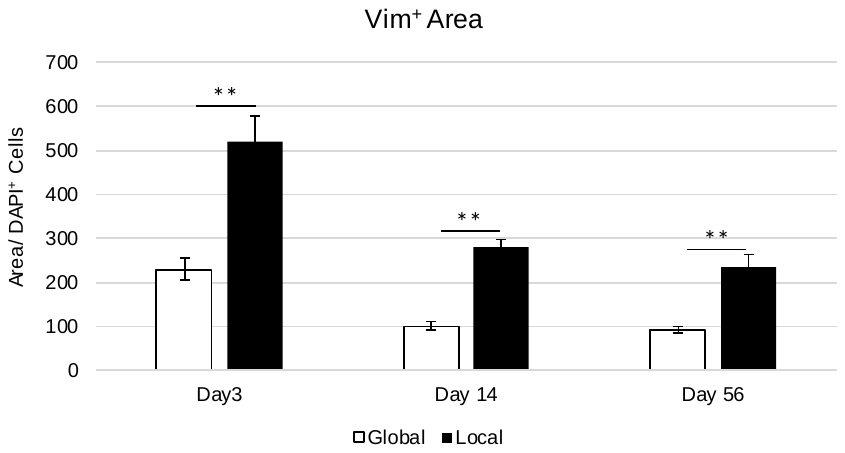

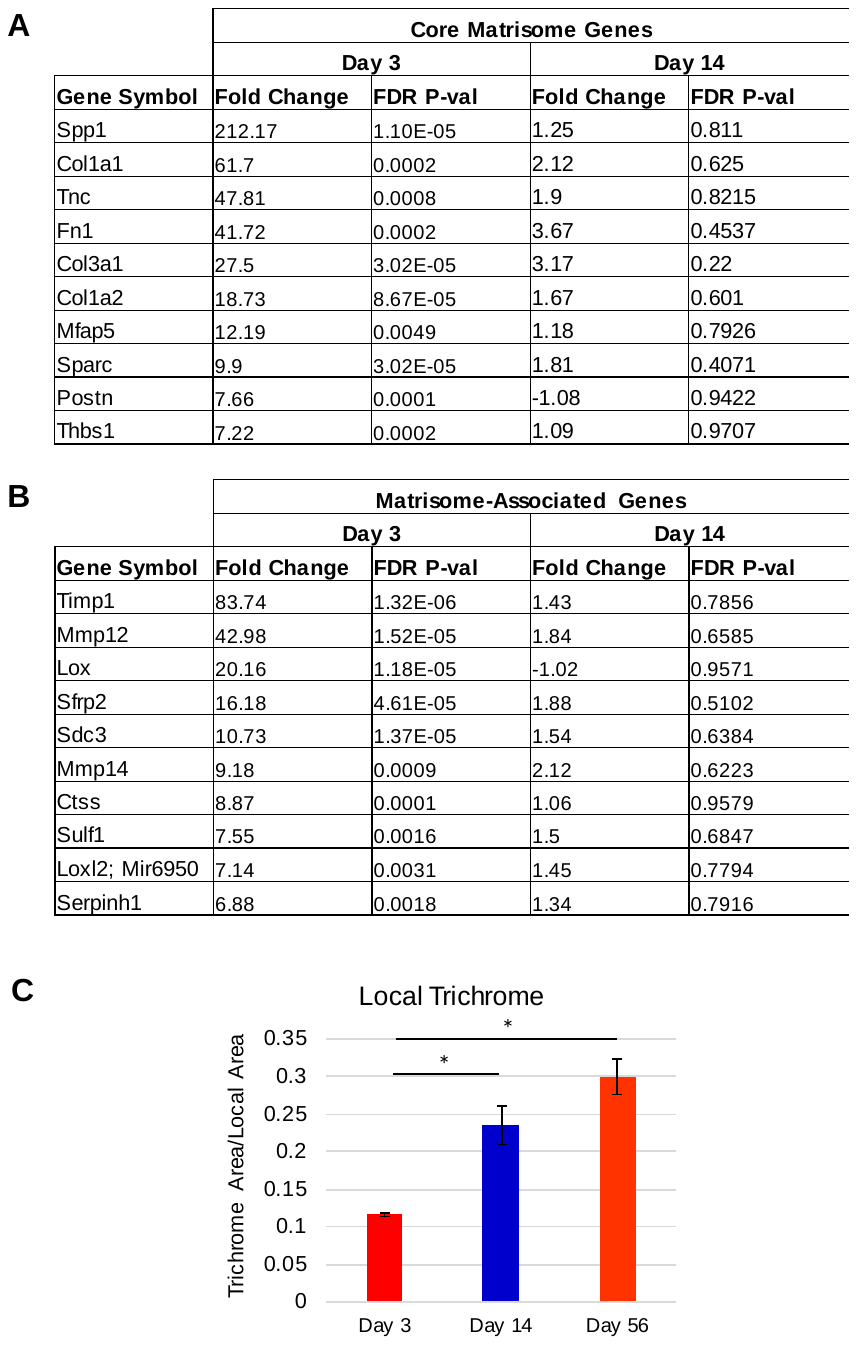

**Appendix Figure 7. Core Matrisome ECM Genes and Matrisome-Associated Genes are Increased 3 Days Post Resection, While Trichrome Increases in the Local Area. A)** At Day 3, the core Matrisome transcriptome is increased, with a return to control levels by Day 14. **B)** Matrisome-associated gene levels also increase at Day 3 but return to control levels by Day 14**. C)** Quantification of the local tissue area covered by blue staining with Masson’s Trichrome staining shows a significant increase in ECM deposition when later time points are compared to the local area at Day 3. The trend suggests that ECM deposition continues to occur in the local area up through 56 days after resection. Day 3, Day 14, and Day 56, n=3. Error bars are S.E.M. Statistical test: One-way ANOVA with Tukey’s HSD post-hoc test. * p≤ 0.05.

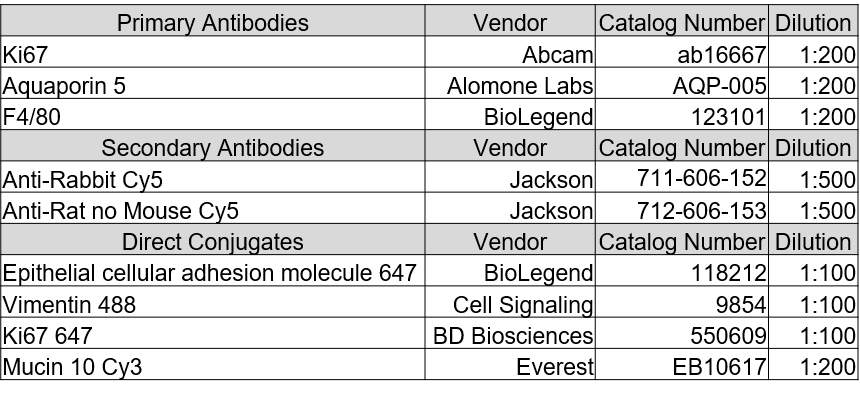

**Table 1: List of Antibodies Used, Manufacturer, Catalog Number and Dilutions.**

**Appendix References:**

Amano O, Mizobe K, Bando Y, Sakiyama K. 2012. Anatomy and Histology of Rodent and Human Major Salivary Glands^|^mdash;Overview of the Japan Salivary Gland Society-Sponsored Workshop^|^mdash; ACTA Histochem Cytochem. 45(5):241–250. doi:10.1267/ahc.12013.

Gao X, Oei MS, Ovitt CE, Sincan M, Melvin JE. 2018. Transcriptional profiling reveals gland-specific differential expression in the three major salivary glands of the adult mouse. Physiol Genomics. 50(4):263–271. doi:10.1152/physiolgenomics.00124.2017.

Lowry R. 2008. VassarStats.

Maruyama C, Monroe M, Hunt J, Buchmann L, Baker O. 2019. Comparing human and mouse salivary glands: A practice guide for salivary researchers. Oral Dis. 25(2):403–415. doi:10.1111/odi.12840.

Nelson DA, Manhardt C, Kamath V, Sui Y, Santamaria-Pang A, Can A, Bello M, Corwin A, Dinn SR, Lazare M, et al. 2013. Quantitative single cell analysis of cell population dynamics during submandibular salivary gland development and differentiation. Biol Open. 2(5):439–447. doi:10.1242/bio.20134309.

Schindelin J, Arganda-Carreras I, Frise E, Kaynig V, Longair M, Pietzsch T, Preibisch S, Rueden C, Saalfeld S, Schmid B, et al. 2012. Fiji - an Open Source platform for biological image analysis. Nat Methods. 9(7). doi:10.1038/nmeth.2019. [accessed 2017 May 22]. http://www.ncbi.nlm.nih.gov/pmc/articles/PMC3855844/.

Zhou Y, Zhou B, Pache L, Chang M, Khodabakhshi AH, Tanaseichuk O, Benner C, Chanda SK. 2019. Metascape provides a biologist-oriented resource for the analysis of systems-level datasets. Nat Commun. 10(1). doi:10.1038/s41467-019-09234-6. [accessed 2019 Jul 23]. http://www.nature.com/articles/s41467-019-09234-6.
